# Developing epidermis acquires nutrients from the external milieu by mTOR-dependent macropinocytosis

**DOI:** 10.1101/2025.03.18.644068

**Authors:** Shravani Reddy Shetty, Arnab Chakraborty, Shivali Dongre, Siddhesh S. Kamat, Mahendra Sonawane

## Abstract

The vertebrate epidermis transitions from a simple bilayer to a complex, multilayered epithelium during development. Periderm, the outermost layer of the bilayered epidermis, is in direct contact with the external aquatic surrounding. While the barrier function of the epidermis is well established, the functional repertoire of the developing epidermis remains unclear. Using zebrafish, we uncover a function of the periderm in nutrient acquisition. Our findings reveal a developmentally regulated wave of macropinocytosis, which is further amplified by extracellular nutrients, leading to the formation of large endo-lysosomal compartments termed Stimulated Nutrient Activated Compartments (SNACs). We identify mTOR as a key integrator of developmental and nutrient-derived signals that modulate SNAC biogenesis. A global lipidomics analysis demonstrates that macropinocytosis-mediated uptake drives the production of metabolically relevant lipids and enhances animal survival when gut function is compromised. Our study identifies a hitherto unappreciated route of nutrient acquisition by the metabolically responsive developing epidermis. We propose that this mechanism fulfils a vital nutritional role, particularly during the transitional phase when maternally provided resources are depleting and the gut is not yet fully operational.

## INTRODUCTION

The epidermis is the outermost epithelial tissue of the integumentary system, which forms a crucial barrier that aids in maintaining the internal milieu of the organism. In mammals, the epidermis originates from a single-layered non-neural ectoderm, which subsequently differentiates into a bilayered tissue comprising an outer periderm and an inner basal layer. As development progresses, the basal layer proliferates and differentiates to generate additional cell strata, whereas the periderm is eventually shed as part of the maturation process (Weiss and Zelickson 1975a; Gerstein 1971; Saathoff et al. 2004; Smart 1970). In contrast, the periderm in fish emerges from an epithelial layer, termed the Enveloping Layer (EVL). Subsequently, the basal layer is established from the non-neural ectoderm following gastrulation, giving rise to an early bilayered epidermis, which eventually stratifies during metamorphosis (Kimmel, Warga, and Schilling 1990; Chang and Hwang 2011).

Composed of flattened squamous cells interconnected by tight junctions, desmosomes, and adherens junctions (Morita et al. 1998; 2002; Sumigray and Lechler 2015; Sonawane et al. 2005), the periderm functions as a provisional barrier. Despite its transient nature, the periderm serves an indispensable role by contributing to the protection and maturation of the developing epidermis and the growing embryo (Weiss and Zelickson 1975b; Wolf 1967). Its aberrant formation has been linked to pathologies such as limb fusion and cleft palate, which result from improper adhesion between adjacent epithelia (Scott et al. 1987; Richardson et al. 2014). The formation of periderm has been described as a “mysterious process in the development of skin” due to its prolonged presence during development and its evolutionary conservation (Wolf 1967). Although traditionally viewed as a transient barrier, whether the periderm has additional functional roles remains unclear.

Ultrastructural analyses of the periderm in developing human foetuses have revealed intriguing features. These include the presence of abundant microvilli, blebs, and both coated and smooth membrane vesicles on its apical surface, which interfaces with the amniotic fluid (Whittaker and Adams 1971; Breathnach 1971; Holbrook and Odland 1975). These observations suggest that peridermal cells actively interact with the external environment, extending the periderm’s role beyond mere protection. Furthermore, the presence of membrane vesicles also raises the possibility that peridermal cells interact with the external surroundings via endocytosis. However, these notions remain to be tested.

Interaction of cells with their surroundings through endocytosis involves the internalization of extracellular substances and plasma membrane components, including receptors. This process supports a multitude of vital functions, such as the modulation of signal transduction pathways, membrane remodelling, immune responses, nutrient uptake, and the regulation of cell division and migration (Grimes et al. 1996; Burke, Schooler, and Wiley 2001; Thottacherry et al. 2018). Endocytosis can be categorized into three main forms: receptor-mediated endocytosis, phagocytosis, and pinocytosis. Among these, macropinocytosis, a form of pinocytosis characterized by non-specific internalization of extracellular fluid via large vesicles called macropinosomes, plays a crucial role in enabling cells to adapt to dynamic external conditions. Although initially recognized for its role in immunity and infection (Norbury et al. 1995; Maréchal et al. 2001), macropinocytosis is now understood as a conserved mechanism across diverse cell types essential for membrane rearrangement, the formation of signalling platforms, and nutrient uptake (Gu et al. 2011; Commisso et al. 2013).

In this study, we uncover a hitherto unknown nutritionally modulated developmental wave of macropinocytosis in the developing periderm. The extent of macropinocytosis and the subsequent formation of vesicles - termed Stimulated Nutrient Activated Compartments (SNACs) - is regulated by mTOR signalling. Internalization of nutrients via macropinocytosis results in a significant increase in key lipid metabolic components, presumably via lipid synthesis, suggesting a critical role of the periderm in nutrient acquisition and possibly in survival under unfavourable conditions.

## RESULTS

### Developing zebrafish epidermis exhibits a developmentally regulated wave of large vesicles originated from macropinocytosis

To understand the presence and extent of the endocytic activity in the peridermal cells, embryos obtained from a transgenic line Tg(cldnB:LynEGFP), which marks the peridermal cell membrane, were pulsed at specific stages either with wheat germ agglutinin (WGA) or fluorophore-conjugated dextran for 4 hours. WGA, which binds to sialic acid residues on the plasma membrane, facilitates tracing of the vesicle membrane, whereas dextran, a marker for fluid-phase endocytosis, labels the lumen of internalized vesicles (Monsigny et al. 1980; Li et al. 2015). Both WGA and Dextran labelled distinct and large endocytic vesicles in the peridermal cells imaged on the head (Figure 1A, S1A). Our quantitative analysis revealed a developmentally regulated wave of endocytic activity, with vesicle numbers peaking at 72 hours post-fertilization (hpf; Figure 1A, 1B). These vesicles were predominantly localized to the apical region of the cell and exhibited an unusually large size of about 0.5-2 μm, which is approximately tenfold larger than typical endocytic vesicles. To determine whether this wave of endocytosis was present in a specific region of the periderm, we extended our analysis to include the flank and fin-fold epithelium. The wave was observed consistently across all examined regions suggesting that this process takes place all over the epidermis rather than being confined to the head peridermal cells (Figure S1B).

**Figure 1:**
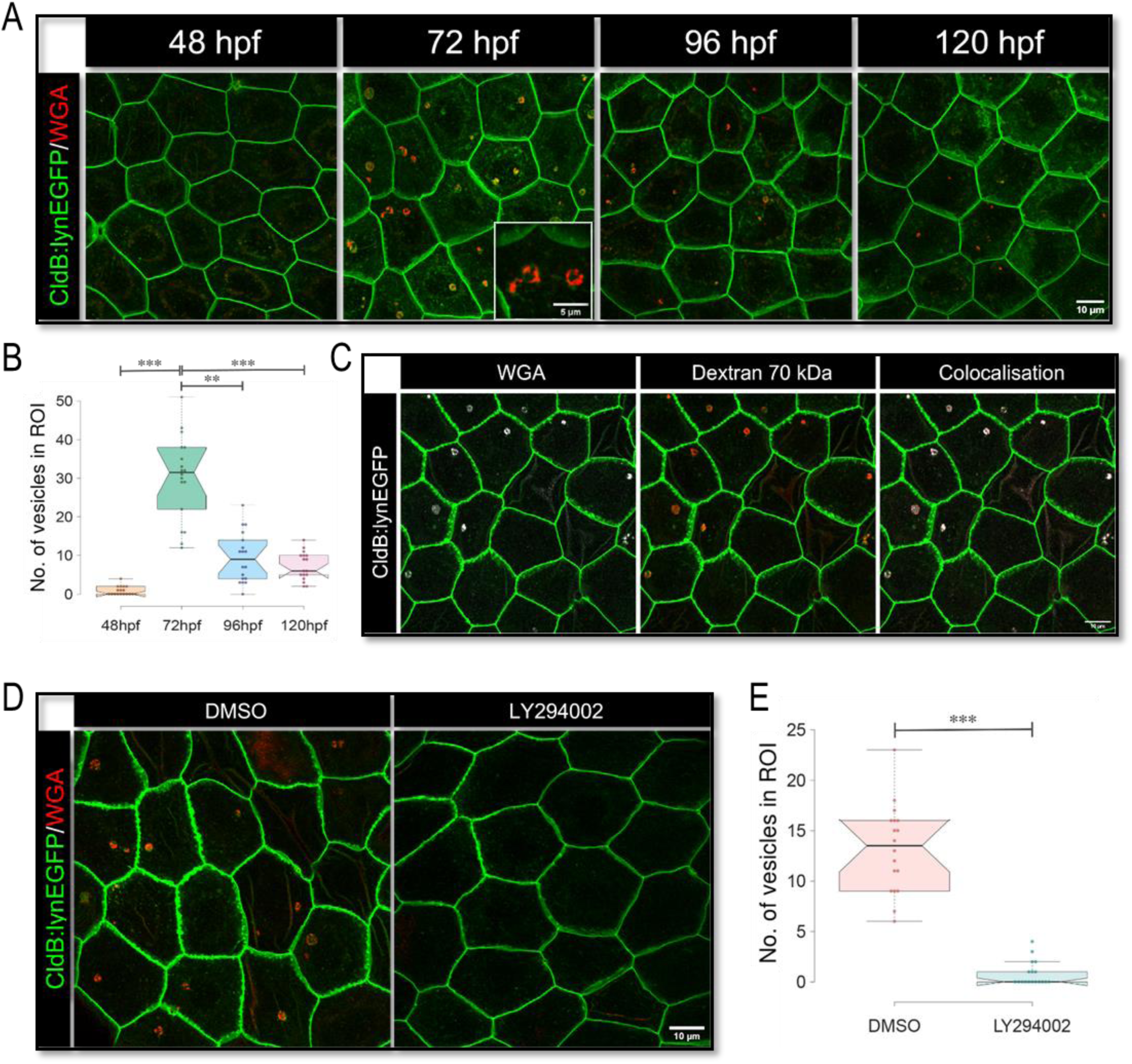
Developing periderm exhibits large endocytic vesicles formed via macropinocytosis in a temporally restricted window. Confocal images of head periderm cells from Tg(CldnB:lynEGFP) embryos (A,C,D), labelled with (A) Lyn-GFP and WGA in a 4-hour live pulse at the mentioned developmental stages in hours post fertilization (hpf), (C) 70kDa dextran and WGA at 72 hpf and (D) WGA in embryos treated with vehicle control (DMSO) and PI3K inhibitor LY294002 (30 μM) for 3 hours. Graphical representation (B,E) of number of vesicles in a given ROI at (B) different developmental stages and (E) upon treatment with LY294002. Vesicle number was quantified from a defined ROI from a total of 16–18 embryos (n) per stage or treatment from three different experimental sets. ***p < 0.001 by one-way ANOVA with Dunn’s post-hoc test in (B) and by two-tailed t-test in (E). hpf = hours post-fertilization; Scale bar = 10 μm.

Given the unusually large size of the vesicles, we examined whether macropinocytosis contributed to their formation by assessing the uptake of high molecular dextran of 70 and 200kDa sizes, which are specifically internalized via macropinocytosis (Li et al. 2015). A live pulse assay at 72 hours post-fertilization (hpf) demonstrated the colocalization of 70 kDa dextran with WGA-labelled vesicles, indicating the involvement of macropinocytosis in vesicle formation (Figure 1C). Additionally, we also observed the presence of 200 kDa dextran in these vesicles (Figure S1C), which further confirmed the ability of periderm to internalize large molecules via macropinocytosis. To further verify that the uptake is macropinocytosis dependent, we inhibited PI3-kinase, a key mediator of the formation of membrane ruffles during macropinocytosis. To achieve this, we treated early larvae at 72 hpf with LY294002 for 3 hrs, with a WGA pulse initiated 1.5 hours after the treatment began (Hoeller et al. 2013). The inhibition of PI3-kinase resulted in a near complete loss of WGA uptake and vesicle labelling in the peridermal cells (Figure 1D, 1E), confirming the macropinocytic contribution to the formation of these vesicles.

To further assess whether the dynamin-dependent microendocytic pathways contribute to these large vesicles, we inhibited dynamin function using dynasore monohydrate. We did not observe any effect on the vesicle number in dynasore treated embryos as compared to the control (Figure S2A, S2B), further corroborating the fact that macropinocytosis is the primary pathway responsible for the formation of these large vesicles.

To conclude, we identified a developmentally regulated wave of apical macropinocytosis culminating in the formation of large vesicular compartments in the entire zebrafish embryonic periderm, with vesicle formation peaking at 72 hpf.

### Peridermal vesicles formed via macropinocytosis are endo-lysosomal in nature

To establish the identity of the macropinocytosis derived large vesicles in the periderm, we asked if they were marked with a particular Rab GTPase. For this, we used transgenic zebrafish lines expressing Rab11-GFP and Rab7-GFP (Clark et al. 2011), which mark recycling endosomes and late endosomes, respectively. We observed that the peridermal vesicles predominantly acquire a late endosomal identity, as evidenced by the presence of Rab7-GFP, but not of Rab11-GFP, on the vesicle membrane (Figure 2A, S2C). We further examined their colocalization with LysoTracker, a marker for acidic compartments, and Lamp1, a lysosomal marker (Pierzyńska-Mach, Janowski, and Dobrucki 2014). Our data revealed that both Lysotracker as well as Lamp1 mark these vesicles, suggesting they are lysosomal in nature (Figure 2B, 2C).

**Figure 2:**
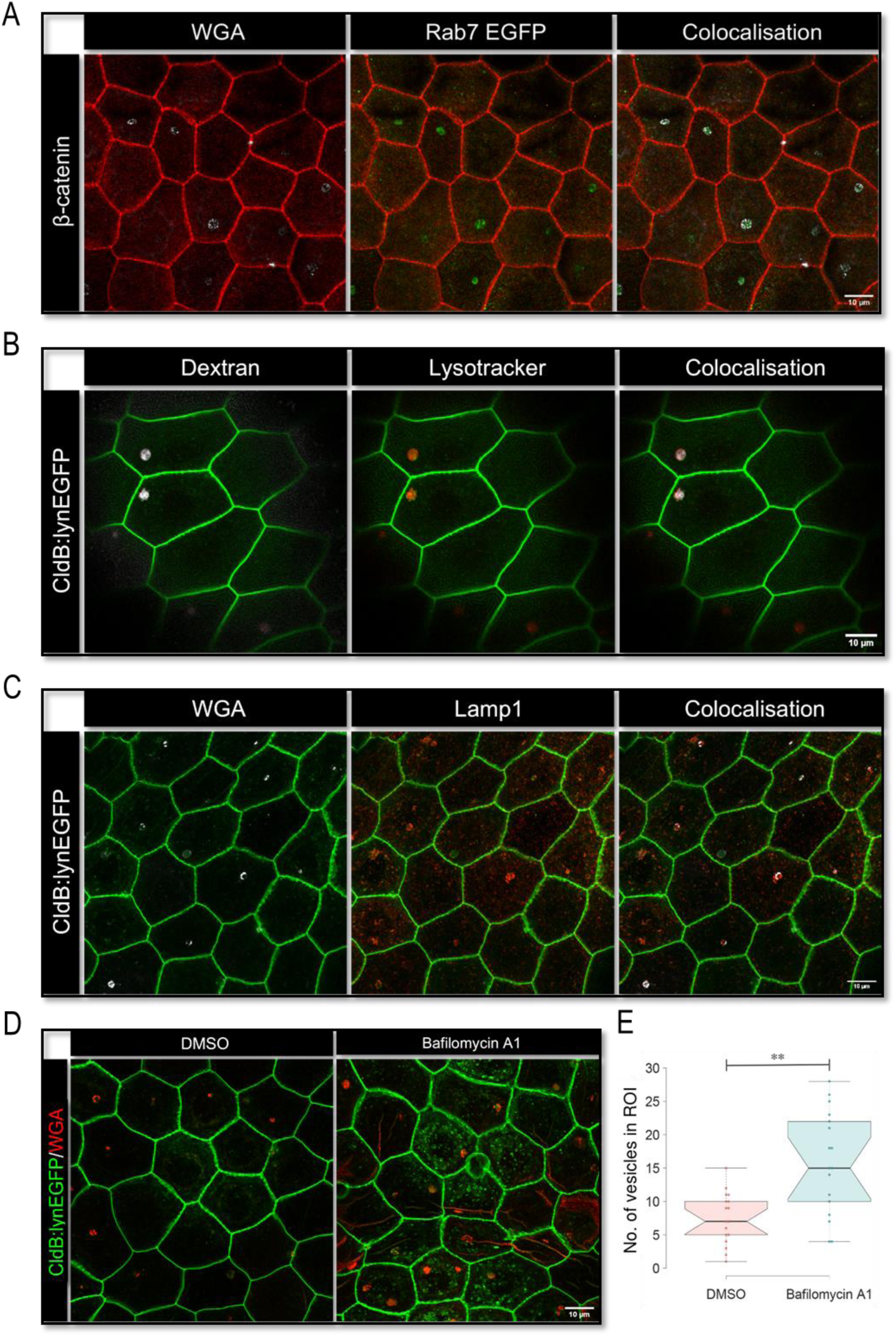
Peridermal endocytic vesicles are endo-lysosomal in nature and possess degradative capacity. Confocal scans of peridermal cells (A-D) marked with β-catenin (A) or lyn-EGFP (B-D) and labelled using (A) 4h WGA pulse in Tg(Rab7:EGFP) embryos at 72 hpf, showing Rab7:EGFP labelled endosomes, (B) Lysotracker and dextran for 4 hours at 72 hours post-fertilization (hpf) and imaged live (C) Lamp1 and WGA at 72 hpf, and (D) 4h WGA pulse after DMSO (vehicle control) and Bafilomycin A1 (40 nM) treatment from 48 to 72 hpf. Notched box-plot (E) showing number of peridermal vesicles per region of interest (ROI) in the DMSO and Bafilomycin A1 treated embryos; a total of 18 embryos (n) were analyzed from three experimental sets. **p < 0.001 by a two-tailed t-test; Scale bar = 10 μm.

To substantiate the degradative function of these compartments, we inhibited lysosomal function by using Bafilomycin A1, which blocks the activity of V-ATPase, an enzyme required for luminal acidification and the activation of lysosomal hydrolases (Wang et al. 2021). The bafilomycin treatment from 48 hpf to 72 hpf resulted in an increased vesicle count as compared to the control (Figure 2D, 2E), suggesting the degradative nature of these vesicles.

In conclusion, the vesicles formed in the periderm via macropinocytosis acquire an endo-lysosomal fate. This indicates a functional role for these compartments in facilitating the breakdown and processing of internalized material, contributing to cellular requirements.

### Nutrient limitation leads to an increase in the number of large endosomal vesicles in the periderm

The role of macropinocytosis in nutrient acquisition to fulfill metabolic requirements is particularly well characterized in cancer cells and organisms such as yeast and Dictyostelium (Williams and Kay 2018; Commisso et al. 2013; Kamphorst et al. 2015). Once internalized, materials are degraded into essential components such as amino acids and lipids, which are essential for sustaining cellular processes (Yoshida et al. 2015).

Zebrafish embryos are lecithotrophic; they rely on maternally deposited yolk for nutrition during early development. However, around 5 days post-fertilization (dpf), the gut becomes functional, and larvae begin to seek food (Kunz-Ramsay 2013; Quinlivan and Farber 2017). To investigate whether nutrient deprivation influences vesicle formation in the periderm, we surgically removed 60-80% of the yolk at 48 hours post-fertilization (hpf). This manipulation led to a qualitative increase in vesicle formation at 72 hpf compared to wounded control larvae, indicating that reduced nutrient availability increases the number of endosomal compartments formed via macropinocytosis in the periderm (Figure 3A-C).

**Figure 3:**
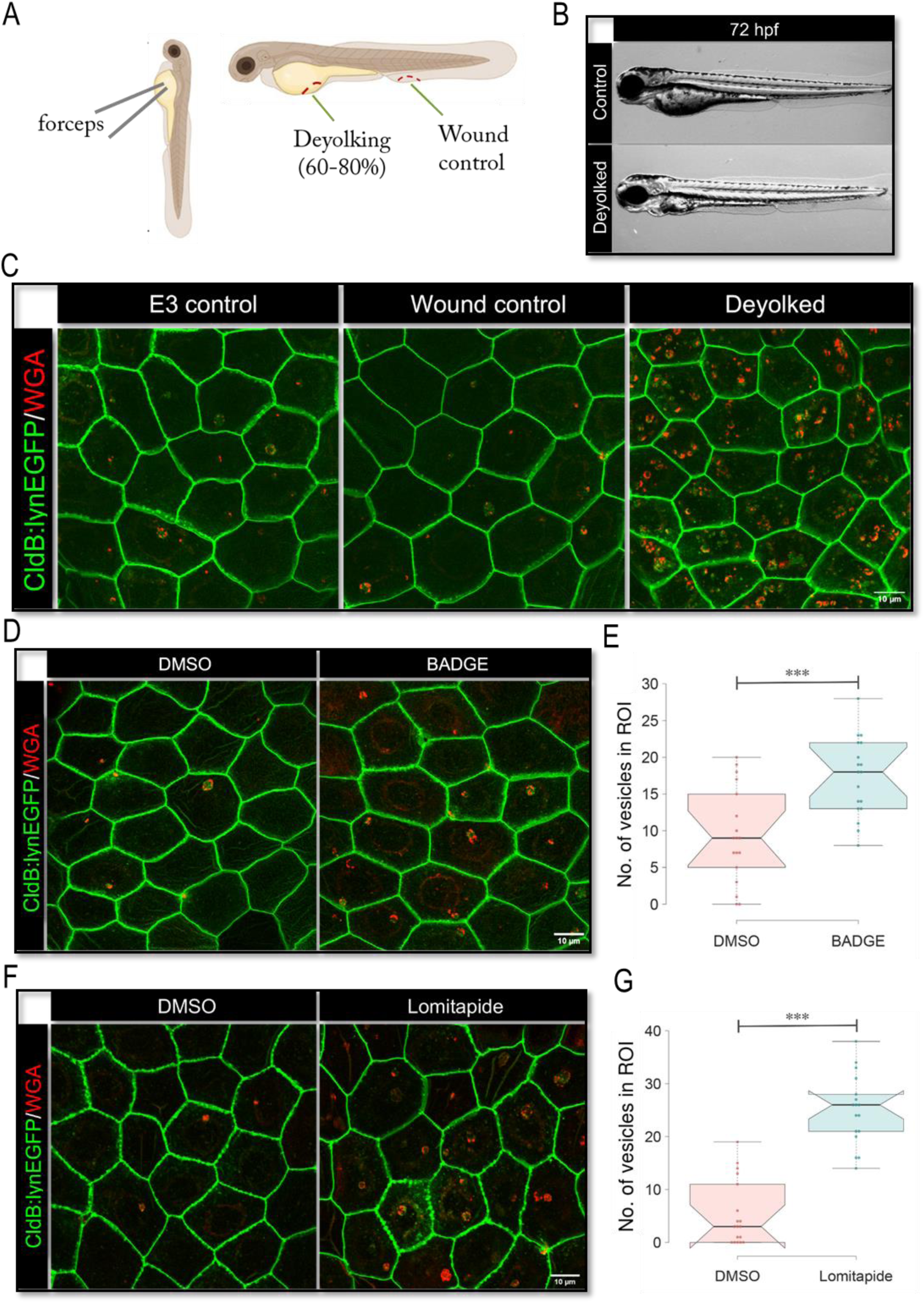
Starvation results in increased vesicle formation in zebrafish periderm. (A) A schematic representation and (B) brightfield images illustrating control and deyolked embryos 24 hours post yolk removal at 72 hpf. Confocal images of peridermal cells labelled with lynEGFP and WGA pulse at 72hpf (C,D,F) in (C) control, wounded, and deyolked embryos, (D) embryos treated with the PPARγ inhibitor BADGE (7.5 μM) for 24 hours starting from 48 hpf and (F) embryos treated with the microsomal triglyceride transfer protein (Mtp) inhibitor Lomitapide (30 μM) for 48 hours starting from 24 hpf. (E,G) Quantification of vesicle numbers within a region of interest (ROI) of the embryos treated with BADGE (E) and Lomitapide (G); n=18 embryos from three experimental sets. ***p < 0.007 by two-tailed t-test; Scale bar = 10 μm.

To further corroborate these findings, we induced starvation using BADGE and Lomitapide, chemical inhibitors of the yolk nutrient transport regulators—PPAR-γ (Fraher et al. 2016) and microsomal triglyceride transport protein (Wilson et al. 2020), respectively. In both these treatments, we observed a similar increase in vesicle formation (Figure 3D-G).

These findings collectively reveal that the organism’s nutritional status plays a pivotal role in the formation of macropinocytosis derived endo-lysosomes within the periderm. As the availability of yolk derived nutrition decreases, macropinocytosis is increased to scavenge nutrients from the extracellular environment. This nutrient-dependent regulation further suggests that macropinocytosis in zebrafish embryos serves as a mechanism to augment nutrient acquisition during nutrient scarcity.

### Presence of complex nutrients in the surrounding medium augments macropinocytosis

As stated earlier, zebrafish embryos rely exclusively on the maternally deposited yolk for nutrition, rather than external sources. Consequently, until feeding commences, embryos and early larvae are typically maintained in an E3 medium, which contains only salts to regulate osmolarity and pH. We hypothesized that as the maternally deposited yolk begins to diminish, the developmental wave of peridermal macropinocytosis facilitates nutrient uptake from the surrounding medium. We, therefore, asked whether adding nutrients to the medium would influence the process of macropinocytosis and the subsequent vesicle formation. To test this, zebrafish embryos at 48 hpf were exposed to an E3 medium supplemented with yolk isolated from 48 h old embryos (see methods for the details) for 24 hours. This stimulation by externally added yolk (referred to as yolk stimulation) resulted in a significant increase in the number of vesicles by 72 hpf (Figure 4A, 4B), demonstrating that the presence of external nutrients enhances macropinocytic activity in the periderm.

**Figure 4:**
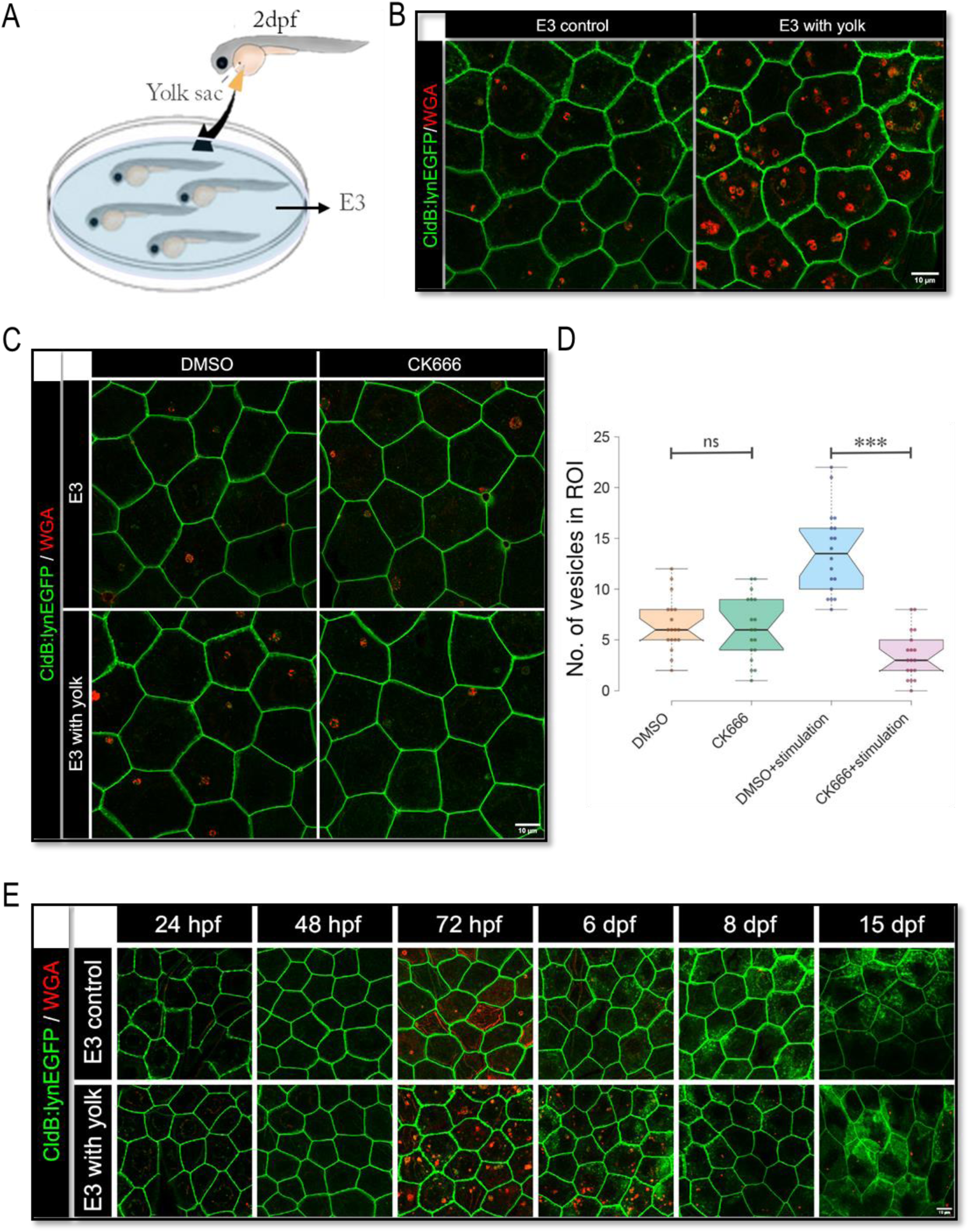
Nutrients in the external milieu stimulate the formation of peridermal vesicles. (A) A schematic illustrating the method of nutrient supplementation by adding yolk to the E3 medium. Confocal images of peridermal cells labelled with lyn-EGFP in embryos maintained in control and yolk-supplemented E3 medium conditions for 24 h followed by (B) WGA pulse at 72 hpf, (C) treatment with the Arp2/3 inhibitor CK666 (150 μM) for 3 hours at 72 hpf, followed by a WGA pulse at 73.5 hpf or (E) after supplementation for 2 h at the various developmental stages mentioned followed by a WGA pulse. Graphical representation (D) of vesicle numbers within a region of interest (ROI) at the conditions mentioned; n=18 embryos from three experimental sets. ***p < 0.0001, ns = non-significant by one-way ANOVA followed by Dunn’s post-hoc test. hpf = hours post-fertilization; dpf = days post-fertilization; Scale bar = 10 μm.

To examine whether diverse biological materials could mediate this effect, embryos were exposed to a range of simple and complex substances, including glucose, glutamine, lipids, BSA, chorion, chicken egg yolk, and zebrafish larval feed. Strikingly, only nutritionally complex, lipoprotein-rich substances such as chorion, egg yolk, and larval feed elicited a significant increase in vesicle numbers (Figure S3A-G). These findings suggest that the induction of vesicle formation is specifically dependent on the presence of lipoproteinaceous components in the medium.

Given that the amplification of the macropinocytic wave depended on complex nutrients, we probed whether the identity of the vesicles was similar to those formed in the absence of externally added nutrients. Utilizing the assays described above, we confirmed that these stimulated vesicles were also derived via macropinocytosis and possess endo-lysosomal fate (Figure S4A-C). Interestingly, CK666 mediated inhibition of Arp2/3 - a regulator of branched actin configuration necessary for membrane ruffle formation and scission of macropinocytic cups - had no effect on vesicle numbers formed during the developmental wave. However, CK666 treatment reduced the number of vesicles upon yolk-stimulation indicating a differential requirement of Arp2/3 for macropinocytosis (Figure 4C, 4D).

We further examined whether nutrient-dependent stimulation is restricted to specific developmental stages. To test this, embryos across various time points, from 24 hpf to 15 dpf, were exposed to yolk for 2 hours. We observed that developmental stages beyond 72 hpf exhibited an increase in vesicle numbers in yolk stimulated embryos as compared to controls, although the magnitude of the increase was smaller than that observed at 72 hpf. Stages earlier than 72 hpf did not exhibit vesicle formation upon yolk exposure, suggesting that yolk stimulation doesn’t override the developmental control (Figure 4E).

To conclude, the exposure of zebrafish embryos to external nutrient, such as yolk, triggers an increase in vesicle number in the periderm, indicating that nutrient availability in the external environment positively modulates macropinocytic activity. While the developmental wave of macropinocytosis diminishes after 72 hpf, this nutrient uptake persists at later stages albeit at a much lower level. Given that this macropinocytosis is nutrient-activated, we named these vesicles as Stimulated Nutrient-Activated Compartments (SNACs).

### mTOR signaling regulates the formation of SNACs in the developing epidermis

The mechanistic target of rapamycin (mTOR) acts as a nutrient and metabolic sensor in eukaryotic cells. Given that the stimulation of macropinocytosis and the formation of SNACs is based on the nutrients available, we asked if mTOR signaling plays a role in regulating the developmental wave of macropinocytosis in the presence of nutrients and their subsequent utilization in peridermal cells.

It is known that mTOR translocates to lysosomes in the presence of abundant amino acids, promoting protein synthesis (Bar-Peled et al. 2012). Immunostaining revealed that mTOR localizes to SNACs in both the control and yolk-stimulated embryos (Figure 5A). Furthermore, embryos exposed to yolk exhibited increased levels of phosphorylated S6 ribosomal protein (pS6), indicating activation of mTORC1 (mTOR complex 1) pathway presumably via lysosomal degradative output (Figure 5B, 5C). To further test whether mTOR signaling regulates the formation of SNACs, we pharmacologically inhibited mTOR function using Torin. We observed a significant reduction in the formation of SNACs under both non-stimulated and nutrient-stimulated conditions upon mTOR inhibition (Figure 5D, 5E), highlighting its essential role in regulating vesicle formation and nutrient uptake within the periderm.

**Figure 5:**
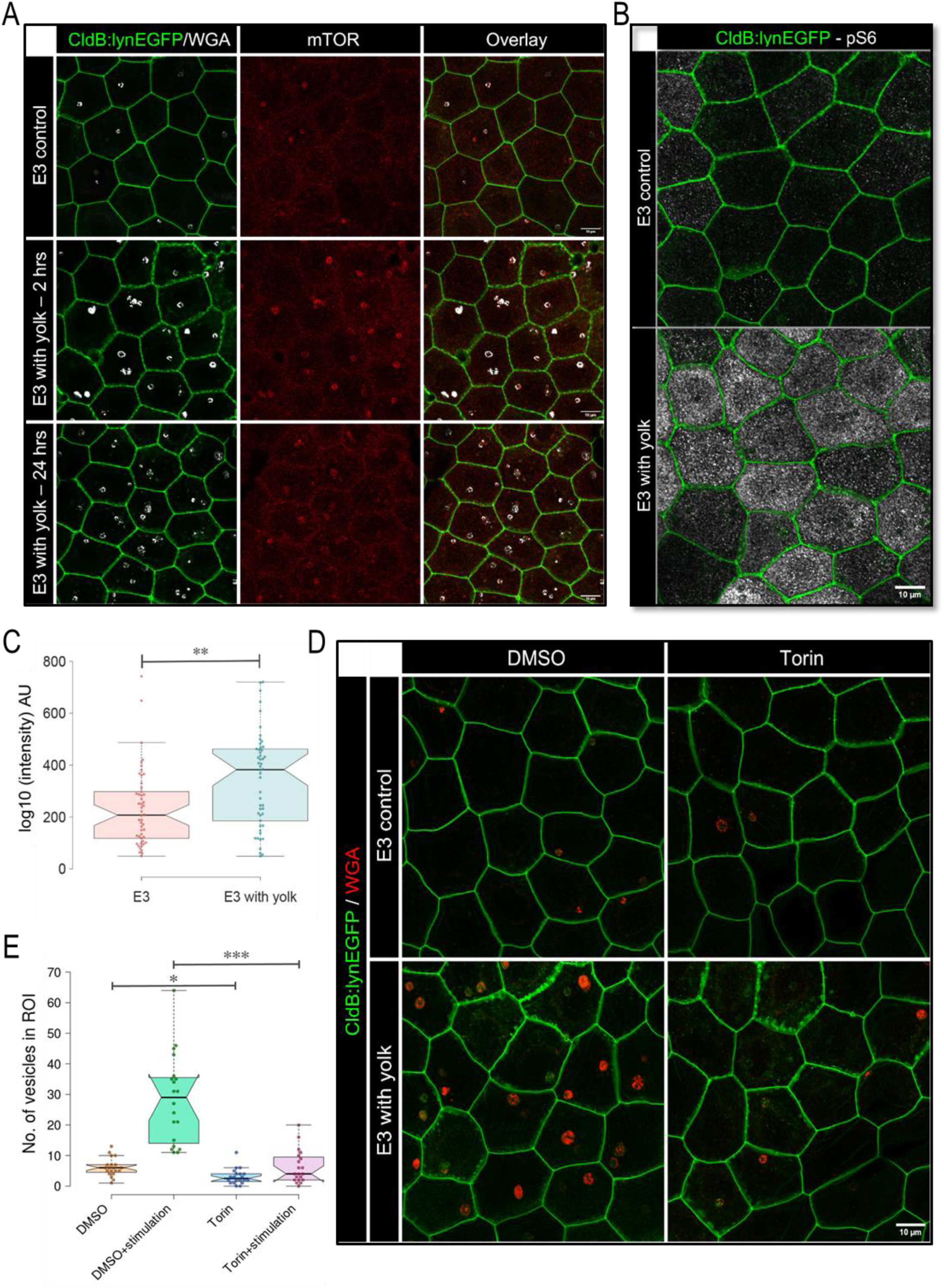
mTOR pathway modulates nutrient-responsive vesicle formation in zebrafish periderm. Confocal scans of the peridermal cells from Tg(CldnB:lynEGFP) embryos reared in control and yolk supplemented E3 medium for 2h (A) or 3h (D) or 24h (A,B) and immune-stained and/or labelled using: (A) mTOR and a 4h pulse of WGA at 72 hours post-fertilization (hpf), (B) phosphorylated S6 (pS6) and (D) 1.5h pulse of WGA at 74.5 hpf after treatment with a vehicle control (DMSO) and mTOR inhibitor Torin (3 μM) for 6 hours at 70 hpf and yolk supplementation at 73 hpf. (C) Quantification of fluorescence intensities in arbitrary units (AU) for pS6; n=50 cells analyzed from 14 embryos from 3 experimental sets, and statistical significance was assessed using a two-tailed t-test, **p = 0.0014. (E) Quantification of vesicle numbers within a region of interest (ROI) after the treatments mentioned, represented in the notched box-plot; 20 embryos were analyzed from three experimental sets, *p<0.01, ***p < 0.0001 by one-way ANOVA followed by Dunn’s post-hoc test; Scale bar = 10 μm.

Our data suggest that upon nutrient exposure, mTOR activation enhances the formation of SNACs via macropinocytosis presumably to facilitate the uptake of available extracellular material.

### Macropinocytosis and SNAC formation promotes lipid synthesis and may aid in embryo growth and survival

Finally, we sought to determine the functional significance of the peridermal macropinocytosis and SNAC formation. To begin with, we evaluated the overall energetic impact of nutrient stimulation on embryos by assessing changes in ATP levels. ATP was extracted from both yolk-stimulated and non-stimulated embryos, which when analyzed revealed a decrease in ATP levels in yolk-stimulated embryos (Figure 6A), suggesting potential energy consumption for the process of macropinocytosis and synthesis of complex biomolecules. To explore this further, we conducted an untargeted lipidomics analysis on whole embryos stimulated with acute (2 h) or chronic (24 h) exposure to yolk, with non-stimulated embryos as normalization controls. Total lipids were extracted from the whole body of embryos at 74 hpf after removing maternally deposited yolk (see Methods for details), followed by an established liquid chromatography coupled mass spectrometry analysis (Kumar et al. 2024; Chakraborty et al. 2025; Jog et al. 2025). The results demonstrated an overall increase in lipid content in yolk-stimulated embryos, with significant elevations in various lipid classes such as diacylglycerols (DAGs), triacylglycerols (TAGs), lysophosphatidylcholines (lyso-PCs), cholesterol, and cholesterol esters (CEs) compared to controls (Figure 6B), all of which are key intermediaries in the biosynthesis of biologically relevant lipids. Additionally, our qPCR analysis revealed upregulated transcription of fatty acid synthase (FASN) after 2 h of yolk-stimulation, which further corroborates the finding that lipid synthesis is enhanced in response to the presence of extra-embryonic nutrients (Figure 6C).

**Figure 6:**
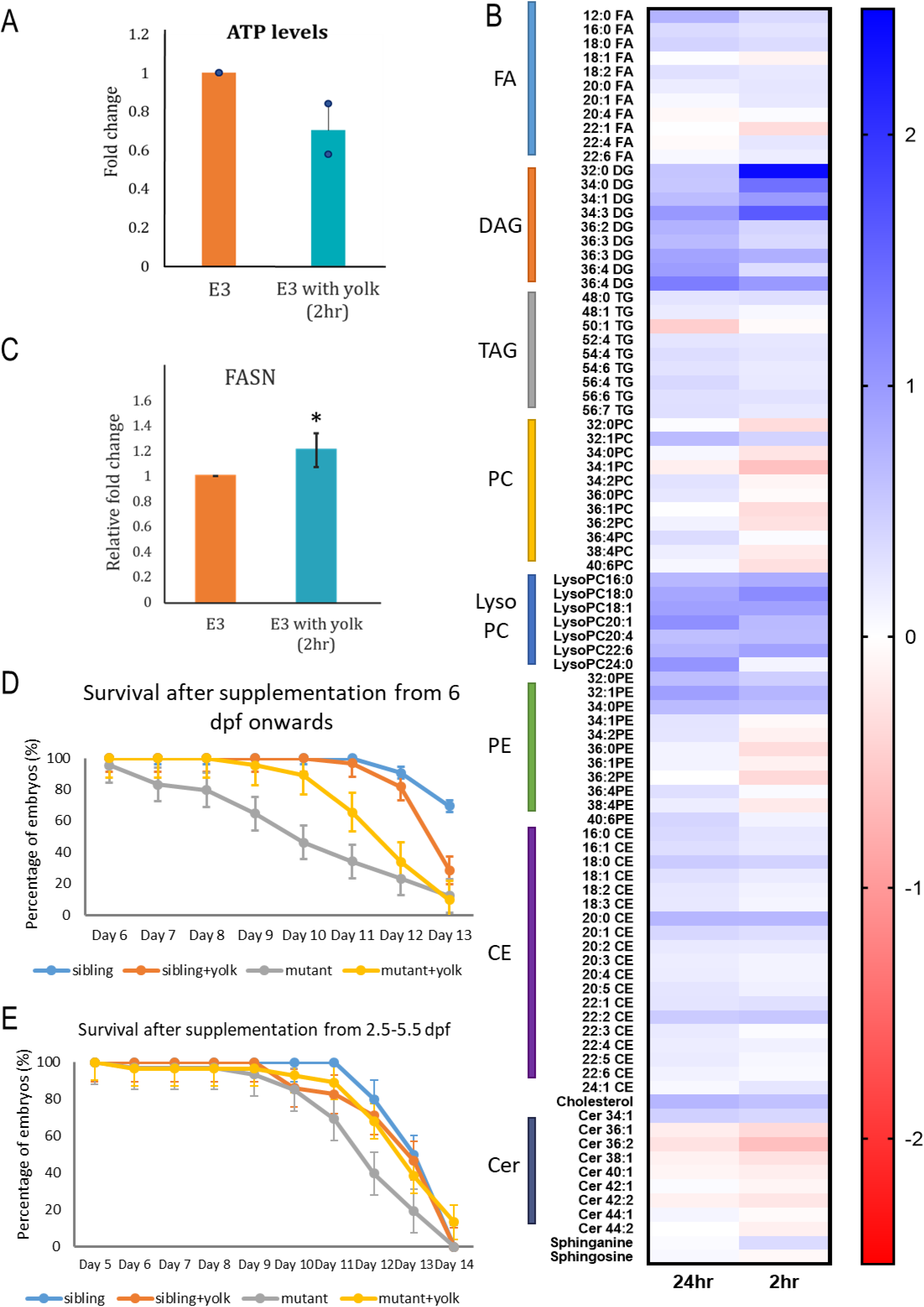
Macropinocytosis promotes lipid synthesis and supports survival in developing zebrafish. (A) The bar graph representing ATP levels in embryos following 2-hour stimulation with yolk-supplemented medium. (B) Heat map of relative log2 fold changes in the lipidome of embryos supplemented with yolk for 2 hours (acute) and 24 hours (chronic) and normalized to controls maintained in E3. Data from 4 replicates; FA = Fatty acids, DAG = Diacylglycerol, TAG = Triacylglycerol, PC = Phosphatidylcholine, PE = Phosphatidylethanolamine, CE = Cholesterol esters, Cer = Ceramides. (C) A bar graph depicting relative fold change in *fasn* gene expression as per qPCR analysis following a 2-hour stimulation with zebrafish embryo yolk. The expression was normalized to *eef1a1*. 20 embryos were pooled to prepare cDNA and 3 biological replicates were considered. *p = 0.02* by Student’s *t*-test. (D,E) Survival graph of *gsp/myosin Vb* sibling and mutant embryos, with or without yolk supplementation (D) from 6 dpf onwards and (E) between 2.5–5.5 dpf. n= 29 for (D) and n=31-34 for (E) from 3 replicates. Data is presented as mean ± SEM.

To evaluate whether macropinocytic uptake supports the organism’s nutritional requirements, we utilized *gsp/myosin Vb* mutants, which exhibit impaired nutrient absorption due to the absence of intestinal folds and reduced microvilli, resulting in mortality by 10–12 dpf (Sidhaye et al., 2016). We reasoned that the peridermal uptake of nutrients via macropinocytosis would compensate, at least partially, for intestinal deficiencies and enhance survival in *gsp/myosin Vb* mutant larvae. We provided yolk supplementation in E3 medium to mutant and control sibling larvae from 6 dpf onwards, when the intestine in wild-type animals is fully functional. Indeed, yolk supplementation significantly improved the survival of *gsp* mutants compared to un-supplemented controls (Figure 6D). To exclude the possibility that the residual nutrient absorption by mutant enterocytes (Sidhaye et al. 2016) contributed to better survival, we performed this assay in an earlier developmental window (2.5–5.5 dpf), a period when the gut is known to be barely functional in zebrafish (Kunz-Ramsay 2013; Quinlivan and Farber 2017). Mutant larvae supplemented with yolk in this early time window again showed significantly improved survival compared to un-supplemented controls (Figure 6E). Together, these results suggest that nutrient uptake via macropinocytosis in the periderm partially compensates for gut impairment, contributing to nutritional requirements during early development.

In conclusion, nutrient-stimulated macropinocytosis in the periderm results in enhanced lipid synthesis. This nutrient-acquisition route may provide an alternative means to support growth and survival, especially under nutrient-limited conditions.

## DISCUSSION

In this study, we establish a previously unrecognized role of the developing periderm in nutrient acquisition through macropinocytosis. Our findings reveal that the periderm actively modulates its macropinocytic activity in response to nutrient availability, thereby facilitating metabolic adaptation. This discovery highlights the functional versatility of the periderm and challenges the prevailing view of it being merely a protective barrier during embryogenesis (Sumigray and Lechler, 2015; Sonawane et al., 2005).

Previous studies in human periderm have reported the presence of vesicular structures (Holbrook and Odland, 1975; Wolf, 1967), yet their origin, composition, and functional relevance has remained elusive. Our investigation identifies a distinct population of large, apically localized vesicles with late endo-lysosomal characteristics in the zebrafish periderm. These vesicles, formed via macropinocytosis, exhibit high sensitivity to nutrient fluctuations, prompting us to designate them as Stimulated Nutrient-Activated Compartments (SNACs). SNACs likely function as dynamic hubs for the degradation and recycling of internalized extracellular nutrients, further facilitating the biosynthesis of critical bioactive lipids essential for development.

We observed that the macropinocytic wave culminating in SNAC formation peaks at 72 hours post-fertilization (hpf). While its magnitude is significantly augmented under nutrient-enriched conditions, the initiation of this process remains developmentally timed. This temporal specificity raises two fundamental questions: what governs the developmental window of macropinocytosis, and how is SNAC formation modulated by the nutrient availability? The emergence of vesicles in the absence of exogenous nutrients suggests that intrinsic developmental cues regulate their baseline formation. Conversely, their amplification under nutrient depletion (yolk removal) or supplementation (external yolk addition) indicates the presence of nutrient-sensing mechanisms that modulate macropinocytic activity.

Our data implicate mTOR signalling as a critical regulator of this process given that pharmacological inhibition of mTOR significantly suppresses both the baseline (developmentally driven) and yolk-stimulated macropinocytosis and SNAC formation. We propose that the developmental wave of macropinocytosis, which requires basal mTOR activity, is a consequence of the depletion of maternally deposited yolk reserves resulting in nutrient scarcity coupled with developmental signals. This plausibly serves as a low-energy consuming mechanism for initial sampling of the external environment. This nutrient sensing, internalisation and hydrolysis to bioavailable substrates such as amino acids, results in the activation of mTOR signalling. This activation likely establishes a positive feedback loop by expending the necessary energy to enhance nutrient acquisition via reinforcing macropinocytosis and SNAC biogenesis through downstream signalling pathways. Thus, mTOR functions as a node that integrates the developmental signal and nutritional status to fine-tune this process. At this stage, it is not clear whether increased macropinocytosis upon yolk depletion and inhibition of yolk utilization is mTOR activation dependent or whether it requires just basal mTOR activity. Further experiments will be required to ascertain this aspect.

The peridermal macropinocytosis that we identified in zebrafish differs from that reported in Xenopus and Hydra vulgaris. In Xenopus, macropinocytosis is induced as an adaptive response to mechanical stress arising from tissue crowding during epidermal proliferation (Bresteau et al., 2024) whereas the surface epithelium of Hydra exhibits constitutive macropinocytosis which is suppressed by high tissue tension (Skokan et al. 2024). However, the zebrafish periderm undergoes this process during a stable, non-proliferative phase at 72 hpf (Sonal et al., 2014; Chouhan et al., 2024). Notably, our AFM data also indicates a progressive increase in tissue tension from 48 to 72 hpf, yet macropinocytosis remains sustained. Therefore, it is unlikely that tissue crowding or mechanical forces dictate this process in zebrafish periderm. Whether macropinocytosis in Xenopus is similarly repurposed for nutrient acquisition remains unknown.

The functional significance of SNACs extends beyond nutrient uptake to metabolic regulation. Indeed, the activation of pS6 downstream of mTORC1 underscores the role of SNAC-mediated nutrient uptake in lipid and protein synthesis. Further, our qPCR analysis revealed a significant upregulation of fatty acid synthase (FASN) following nutrient stimulation. Consistently, the lipidomics analysis demonstrated elevated levels of key lipid intermediates, including DAGs, TAGs, lyso-PCs, cholesterol, and cholesterol esters, in nutrient-stimulated embryos. This is interesting given that phosphatidylcholine has been implicated in channeling newly synthesized fatty acids into membrane biogenesis under pro-growth conditions (Quinn et al., 2017). Besides, TAG synthesis is known to play a protective role by mitigating lipotoxicity during fatty acid overload (Piccolis et al., 2019) and prevent mitochondrial dysfunction under nutrient deprivation (Nguyen et al., 2017). Furthermore, studies have also shown that TAG synthesis redirects fatty acids away from pathways that produce ceramides and other stress-associated lipids (Ackerman et al. 2018). Consistently, our lipidomics data show a decrease in ceramide levels, suggesting a protective mechanism activated by SNAC induction to mitigate lipid-induced stress.

The physiological significance of SNAC-mediated nutrient acquisition is highlighted by the improved survival of *gsp/myosin Vb* mutant zebrafish larvae having an impaired gut function. Although the possibility of partial gut functionality in these mutant larvae presents a caveat to the survival assay, the observed survival advantage, together with lipid remodelling following nutrient uptake via SNACs, suggests that peridermal macropinocytosis can transiently compensate for intestinal deficiencies. It further indicates that the nutrients acquired via the peridermal route contribute systemically to sustain metabolism and promote survival under starvation conditions. Further studies are necessary to validate this notion and to elucidate the mechanisms by which periderm-derived nutrients are transported systemically.

Initially evolved as a nutrient acquisition mechanism in unicellular organisms like yeast and Dictyostelium, macropinocytosis was later adapted by nutrient-deprived cancer cells as a critical means to sustain metabolic demands (Lin, Mintern, and Gleeson 2020). There is sporadic evidence of the conservation of this process in specialized mammalian cells, such as lysosome-rich enterocytes in the neonatal gut epithelium (Park et al., 2019) and trophoblasts in the foetal environment (Shao et al., 2021). However, its broader significance during vertebrate development has remained largely unexplored. Our work reveals macropinocytosis as a conserved mechanism operating in a normal developmental context to acquire extracellular nutrients.

Collectively, this study establishes macropinocytosis in the zebrafish periderm as an essential developmental adaptation that links environmental nutrient availability to intracellular metabolic pathways via mTOR activation. By facilitating nutrient acquisition, degradation, and metabolic reprogramming, the periderm emerges as a metabolically dynamic tissue critical for survival in resource-limited conditions. These findings redefine the functional repertoire of the periderm and provide a framework for future exploration of macropinocytosis as a developmental strategy bridging environmental constraints and metabolic needs.

## MATERIALS AND METHODS

### Ethics Statement

Zebrafish husbandry and experimental handling were conducted in accordance with the guidelines recommended by the Committee for the Purpose of Control and Supervision of Experiments on Animals (CPCSEA), Government of India, and approved by the Institutional Animal Ethics Committee (TIFR/IAEC/2019-6 and TIFR/IAEC/2024-5).

### Fish Strains

All experiments were carried out using the wildtype Tübingen (Tü) strain. Transgenic lines Tg(CldB:LynEGFP) (Haas and Gilmour, 2006), Tg(h2afx:EGFP-Rab7)mw7, and Tg(h2afx:EGFP-Rab11a)mw6 (Clark et al., 2011) were utilized to mark the plasma membrane, late endosomes, and recycling endosomes, respectively. For the survival assay, *goosepimples* mutant allele *gsp^K38b^* was used (Sidhaye et al. 2016).

### WGA, Dextran, and LysoTracker Assays

To assay endocytosis in live periderm, embryos were incubated with 5 μg/ml WGA (Invitrogen, W11262, W32464, W32466), 50 μg/ml Dextran (Invitrogen D-22911, D-1818, D-7139), or 5 μM LysoTracker Red DND-99 (Invitrogen, L7528) in 1X E3 without methylene blue for 4 hours at 28.5°C unless specified otherwise. Following incubation, embryos were washed three times with E3 for 15 minutes each to remove excess dye. Embryos were then either imaged live or fixed in 4% paraformaldehyde (PFA) in PEMTT buffer (0.1 M PIPES, 5 mM EGTA, 2 mM MgCl₂, 0.1% Triton X-100, 0.1% Tween 20, pH 6.8) overnight (Song et al. 2013). Fixed samples were subjected to a glycerol upgrade (30%, 50%, and 70%) before imaging. Approximately 20% of embryo clutches displayed little to no vesicle formation at 72 hpf; these clutches were excluded from the analysis.

### Whole-Mount Immunostaining

For visualization and quantification of specific proteins in the periderm, embryos were fixed in 4% PFA in PEMTT buffer for 30 minutes at room temperature, followed by overnight incubation at 4°C. For mTOR staining, an additional permeabilization step was performed with a methanol series (30%, 50%, 70%, and 100%) followed by overnight incubation in absolute methanol at -20°C. For immunostaining, embryos were serially rehydrated, followed by 5 washes with PBT (0.8% Triton X-100 in PBS) of 10 min each and then blocked in 10% normal goat serum (NGS; Jackson ImmunoResearch Labs, 005-000-121) in PBT for 4 hours. Primary antibodies were incubated overnight at 4°C with 1% NGS in PBT using the following dilutions: mouse β-catenin (1:400; Sigma, C7207), mouse Lamp1 (1:100; ABclonal, A16894), rabbit pS6 (Ser240/244) (1:100; CST, 2215S), rabbit mTOR (1:100; CST, 2893S), and chicken anti-GFP (1:200; Abcam, ab13970). Unbound antibodies were removed by five 30-minute PBT washes, followed by incubation with Cy3-conjugated secondary antibodies (1:750; Jackson ImmunoResearch Labs, 115-165-144/146) for 3–4 hours at room temperature. After six PBT washes of 15 minutes each, embryos were post-fixed in 4% PFA in PEMTT, subjected to a serial glycerol upgrade, and mounted in 70% glycerol for imaging.

### Deyolking, Wound Control, and Stimulation Assays

At 48 hpf, embryos were dechorionated and transferred to a petri dish with minimal E3 medium. For deyolking, a small incision was made in the yolk sac using fine-tip forceps (Dumont, INOX #5), and using the blunt end of the forceps, gentle pressure was applied to release 60–80% of the yolk. Approximately 12–14 deyolked embryos were placed in fresh E3 medium. After 24 hours, embryos showing effect of injury, such as heart edema, were excluded. Wound control embryos were generated by inflicting a minor injury on the finfold adjacent to the yolk extension using fine-tip forceps.

Yolk stimulation assays were performed by isolating yolk from 48 hpf embryos. Approximately 80–90% of the yolk was extracted by creating a small tear in the yolk region of the embryo using fine-tip forceps. Once the yolk began to flow due to the external pressure difference, further extraction was facilitated by gently pressing on the yolk, allowing it to mix with 1–2 drops of E3 medium surrounding the embryo. After removing the embryo body with forceps, the yolk-E3 mixture was pipetted up and down using a Pasteur pipette to ensure complete dispersal of yolk granules into the E3 medium and to prevent adhesion to the glass slide. The resulting yolk-E3 suspension was then transferred to a Petri dish containing fresh E3 medium, which was subsequently used for stimulation by introducing the embryos. Typically, yolk extracted from eight embryos was suspended in 4 mL of E3 medium in a 30-mm Petri dish, accommodating 12–15 embryos for stimulation. The ratio of yolk to E3 medium volume, dish size, and embryo number was proportionately adjusted as needed. Yolk stimulation was conducted for either 2 hours or 24 hours to assess acute and chronic effects, respectively. Additional stimulants included 1% egg yolk in E3, 2.5 mg/ml larval feed in E3, 0.5% BSA, 75 mM glutamine, 30 mM glucose, or 10 μg/ml lipid mixture, all administered for 24 hours. Variability in vesicle numbers following 24-hour embryo yolk stimulation was observed across clutches and experiments. Notably, 20% of these sets exhibited either no increase or small, dispersed vesicles rather than the large vesicles reported in the results. These sets were excluded from the analysis.

### Drug Treatments

Drug stocks were prepared in DMSO (Sigma-aldrich, D8418). All treatments were conducted in 1% DMSO in E3 for both control and test conditions. LY294002 (Abcam, ab120243) and CK666 (Calbiochem, 182515) were used at 30 μM and 150 μM concentrations, respectively, to treat embryos at 72 hpf for 3 hours. Treatments of Dynasore Monohydrate (30 μM) (Sigma, D7693) and Bafilomycin A1 40 (nM) (Sigma, 1793) were done at 48 hpf for 24 hours. BADGE (7.5 μM) (Tocris, 1326) and Lomitapide (30 μM) (Caymann, 10009610) treatments began at 24 hpf; the inhibitors were refreshed at 48 hpf. Torin (5 μM) (Abcam, ab218606) was applied at 70 hpf for 6 hours. After the treatments are over, embryos were washed three times in E3 before fixation.

### ATP Assay

Cellular ATP was extracted using a boiling water method. Five embryos per condition were transferred to microcentrifuge tubes, E3 medium was removed, and 1 mL of ATPase-free boiling water (provided in the ENLITEN ATP kit, Promega, FF2000) was added. After heating at 95°C for 5 minutes, embryos were homogenized with a pestle and reheated. The homogenate was centrifuged at 12,000 g for 15 minutes at 4°C, and 20 µL of the supernatant was used for bioluminescence measurement using the ENLITEN ATP kit, with ATP standard curves prepared by serial dilutions of 1 μM ATP. ATP values were normalized to protein content via a Bradford assay.

### Lipidomics

Samples comprising 70 embryos per condition (control, acute 2-hour stimulation, chronic 24-hour stimulation) were processed. At 74 hpf, embryos were washed three times with E3 and treated with Ginsburg fish Ringer’s solution without Ca^2+^ (111.2mM NaCl, 3.35mM KCl, 2.38mM NaHCO_3_) to dissolve and remove yolk through gentle pipetting and thermomixing (1,100 rpm, 7 minutes, 16°C). The yolk was separated via centrifugation (5,000 g, 5 minutes, 4°C). Embryos were washed in PBS, snap-frozen in liquid nitrogen, and stored at -80°C until further processing.

The embryos were resuspended in 620 µl of chilled PBS and were homogenized on an ice bed using a sonicator for 1 min (2 sec on and 4 sec off). 10 µl of this homogenate from each sample was collected for protein estimation required for data normalization during lipid analysis. The rest of the sample was used for extraction using a modified Folch Method previously reported (Kelkar et al. 2019; Sen et al. 2023). Briefly, the homogenates were made up to 1mL with PBS, followed by the addition of Chloroform (CHCl_3_) and methanol at the ratio of 2:1:1 (CHCl_3_: Methanol: PBS). For semi-quantitative analysis of lipids, 1 nmol of C15:0 MAG (Synthesized as reported previously (Joshi et al. 2018)) and 1 nmol of pentadecanoic acid (C15:0 FFA) (Sigma Aldrich, Catalog # P6125) were added as internal standards for positive and negative mode analytes respectively.

This homogenate mixture was vigorously vortexed and centrifuged at 3000 x g for 15 minutes to separate the mixture into an organic phase (bottom) and an aqueous phase (top) separated by a protein disk. The organic phase was removed by pipetting and stored on ice (at ∼ 0 – 4 °C), while 100 μL of formic acid (MS grade, Honeywell, Catalog # 94318) was added to the aqueous phase to enhance the extraction of phospholipids. This mixture was vigorously vortexed, and 2 mL of CHCl3 was added, after which the mixture was again mixed and centrifuged at the same conditions. The organic layer from this extraction was pooled with the previously obtained organic layer and dried under a stream of nitrogen gas. Protein estimation was performed using the Pierce BCA assay (Thermo).

The dried lipid extracts were resolubilized in 200 μL of 2:1 CHCl3: MeOH and 10 μL was injected into an Agilent 6545 Quadrupole Time-Of-Flight (QTOF) LC-MS/MS for semi-quantitative analysis using high-resolution auto MS/MS methods and chromatography techniques described previously (Chakraborty et al. 2025; Jog et al. 2025). In brief, a Gemini 5U C-18 column (Phenomenex) coupled with a Gemini guard column (Phenomenex, 4x3 mm, Phenomenex security cartridge) was used for LC separation. The solvents used for the LC-MS analysis were buffer A: 95:5 water: methanol and buffer B: 60:35:5 isopropanol: methanol: water. For negative ion mode analysis, 0.1% (v/v) ammonium hydroxide was added in each buffer, while 0.1% (v/v) formic acid + 10 mM ammonium formate was used as additives for analysis in the positive ion mode. Methods spanned 60 minutes, starting with 0.3 mL/min 100% buffer A for 5 minutes, 0.5 mL/min linear gradient to 100% buffer B over 40 minutes, 0.5 mL/min 100% buffer B for 10 minutes, and equilibration with 0.5 mL/min 100% buffer A for 5 minutes. All LC-MS runs were performed using an ESI source with the following MS parameters: drying and sheath gas temperature = 320 °C; drying and sheath gas flow rate = 10L/min; fragmentor voltage = 150 V; capillary voltage = 4 kV; nebulizer (ion source gas) pressure = 45 Ψ and nozzle voltage = 1 kV. For the analysis of different lipids, a curated lipid library was employed in the form of a Personal Compound Database Library (PCDL), and the peaks were validated based on relative retention times and MS/MS fragments obtained. Quantifying all lipid species involved normalizing areas under the curve to the corresponding internal standard area and further normalizing to the protein concentration.

### Survival Assay

Survival assays were performed using *gsp^K38b^* mutant fish. Embryos were phenotypically sorted at 48 hpf into mutant and sibling control groups, based on the epidermal phenotype, which recovers over time (Sonal et al. 2014). The mutant and wild-type siblings were maintained separately. For nutritional supplementation, 10–12 embryos per condition (mutant+yolk), (sibling+yolk) were exposed to external nutrient supplementation (see Stimulation Assays) during specific developmental windows. For the 2.5–5.5 dpf window, yolk supplementation was administered for 12 hours per day to ensure proper swim bladder development. Embryos were alternately incubated in yolk-supplemented E3 medium and plain E3 medium every 6 hours to optimize exposure with yolk. For a supplement regime starting at 6 dpf onward, larvae were exposed to the yolk for 24 hours but the yolk was replaced daily to avoid its decay. Mortality was recorded daily and the dead larvae were removed from the plate. The survival percentages were calculated as the ratio of live embryos to the initial total. Survival data were plotted as percentage survival over time. Sets exhibiting unnatural deaths due to handling or other factors, including failure of swim bladder development, were excluded. Two sets from the paradigm involving yolk supplementation after 6 dpf and one set from the 2.5–5.5 dpf supplementation paradigm were omitted from the analysis.

### RNA Isolation and Quantitative PCR (qPCR)

RNA was isolated from whole embryos by processing approximately 20 embryos per condition. Embryos were transferred into 1.5 ml microcentrifuge tubes and excess E3 medium was removed. Subsequently, 200 µl of chilled TRIzol (Invitrogen, 15596026) was added, and embryos were homogenized using a pestle while the tubes were kept on ice. An additional 800 µl of TRIzol was added to each tube, followed by overnight storage at - 80°C. After thawing on ice, 200 µl of chloroform was added to each tube, mixed thoroughly, and centrifuged to separate the phases. The RNA-containing aqueous phase was carefully transferred to a fresh microcentrifuge tube, and RNA was precipitated by adding 1 ml of 100% isopropanol and 1 µl of GlycoBlue (Invitrogen, AM9516) for improved visualization. The mixture was incubated overnight at -20°C. Following centrifugation, the RNA pellet was washed with 75% ethanol, air-dried at 45°C, and resuspended in nuclease-free water. RNA concentration and quality were assessed before proceeding with cDNA synthesis.

Complementary DNA (cDNA) was synthesized using the SuperScript IV First-Strand Synthesis kit (Invitrogen, 10891050) as per the manufacturer’s protocol. Quantitative PCR (qPCR) was performed using GoTaq qPCR Master Mix (Promega, A6001). cDNA samples from all conditions were analysed in triplicate, and the average cycle threshold (Ct) values were calculated. Target gene expression (*FASN*) was normalized against the control housekeeping gene *eef1a1*. Primer sequences used for amplification are listed below:

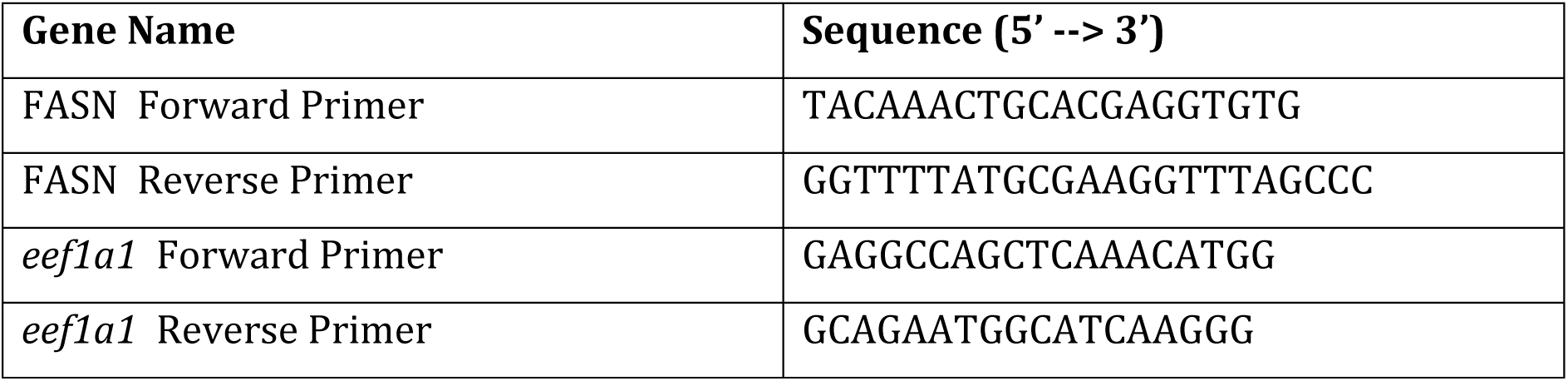

### Image Acquisition and Processing

Epidermis in the dorsal head region was imaged from the immune-stained embryos and larvae unless otherwise specified using a Zeiss LSM 880 confocal microscope with a Plan-APOCHROMAT 63X/1.4 oil objective and a 1.5× optical zoom. Live imaging was conducted with partially submerged anesthetized embryos using a Plan-APOCHROMAT 40×/1.0 water immersion objective. Image processing and analysis were performed using ImageJ (FIJI).

### Image Analysis

Image analysis and processing were performed using ImageJ (FIJI). Vesicles were quantified in an ROI by visual counting; the dimensions of ROI were kept constant for the conditions during imaging. Vesicles having a size lower than 200nM were not counted, to reduce the noise from endosomes and lysosomes present in cells at all developmental stages. The transgenic line, Tg(CldB:LynEGFP) was used to perform pS6 staining and lyn-EGFP was used as a cell boundary marker. The polygon tool was used on composites of pS6 and GFP channel to mark cell boundaries using the GFP channel as a reference. From the pS6 channel, the mean intensity was measured for each slice of the entire Z-stack. The average intensity per cell was calculated and plotted.

### Statistical Analysis

Quantified data were visualized using R. Box-jitter plots displayed medians and interquartile ranges (25th–75th percentile), with error bars indicating 95% confidence intervals. Statistical significance was assessed using Mann-Whitney U tests or Kruskal-Wallis tests with post hoc analysis as appropriate.

## ACKNOWLEDGMENTS

We thank Dr. Kalidas Kohale for maintaining the fish facility and Boby R.V. for overseeing the microscope facility. We also thank Saddam Shekh for the maintenance of the LC-MS facility at IISER Pune. We also acknowledge TIFR-DAE (RTI4003: 1303/2/2019/R&D-II/DAE/2079) to M.S., and the Science and Engineering Research Board (SERB), Department of Science and Technology, Govt. of India (Grant: SB/SJF/2021-22/01) to S.S.K. for funding support.

## AUTHOR’S CONTRIBUTION

Performed experiments - RS, AC; Data analyses and interpretation, Zebrafish experiments - RS, MS; Lipidomics experiments - AC, RS, SSK; Noted the presence of large vesicles in the periderm for the first time - SD; Conceived and conceptualised the project - RS, MS; Project administration and Funding acquisition - MS, SSK.

---

**Supplementary Figure 1:**
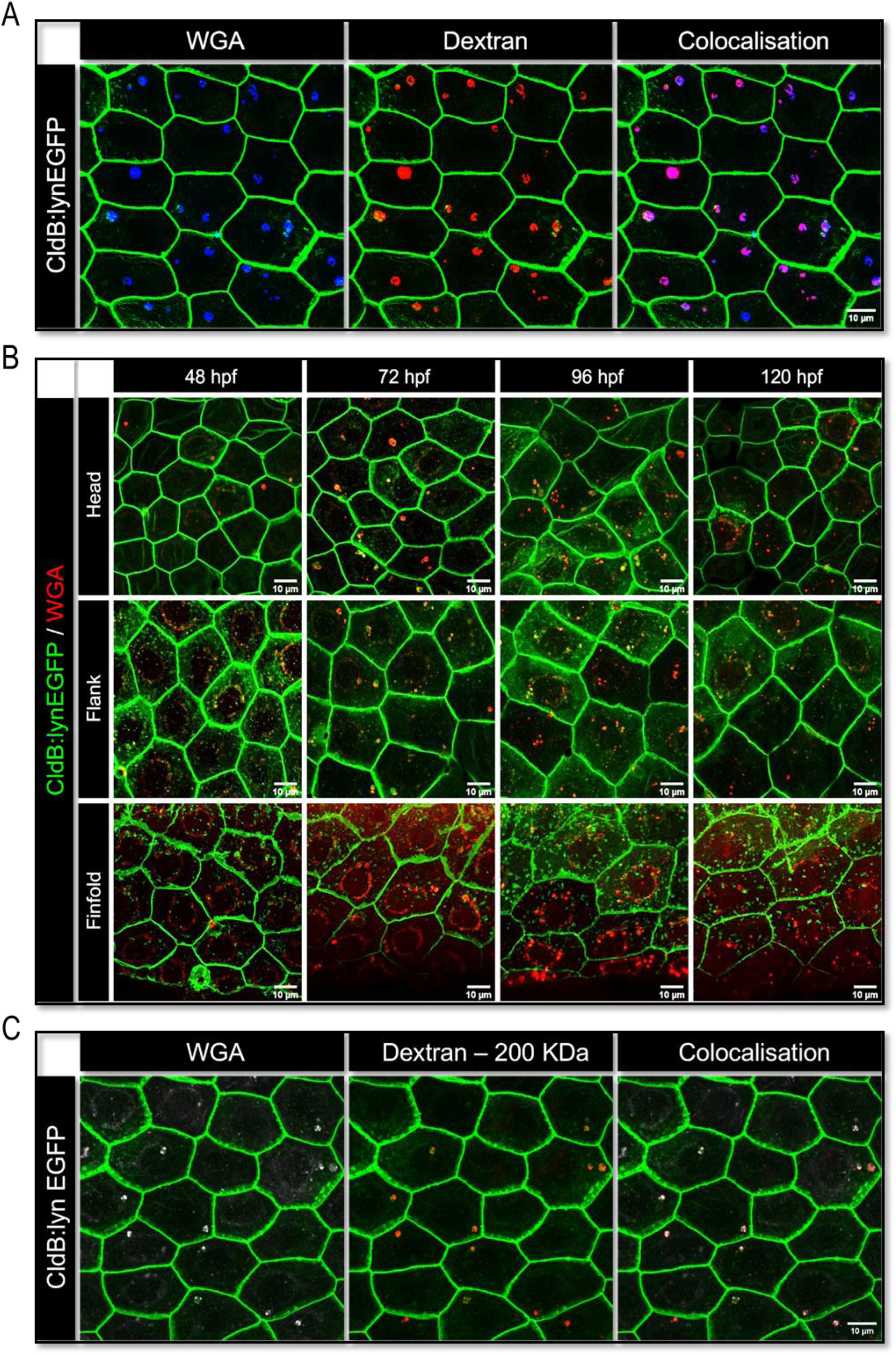
Spatiotemporal distribution of macropinocytic vesicles across periderm. Confocal images of peridermal cells from *Tg(CldnB:lynEGFP)* embryos pulsed with WGA (A-C) showing (A) the uptake of 10 kDa dextran by WGA-labelled vesicles at 72 hpf, (B) WGA-labelled vesicle formation in the peridermal cells covering the head, flank, and finfold during 2 to 5 days post-fertilization (dpf) time-window, and (C) co-localization of 200 kDa dextran with WGA-labelled vesicles at 72 hpf corroborating macropinocytic uptake specificity in peridermal cells. hpf= hours post fertilization; Scale bar = 10 μm.

**Supplementary Figure 2:**
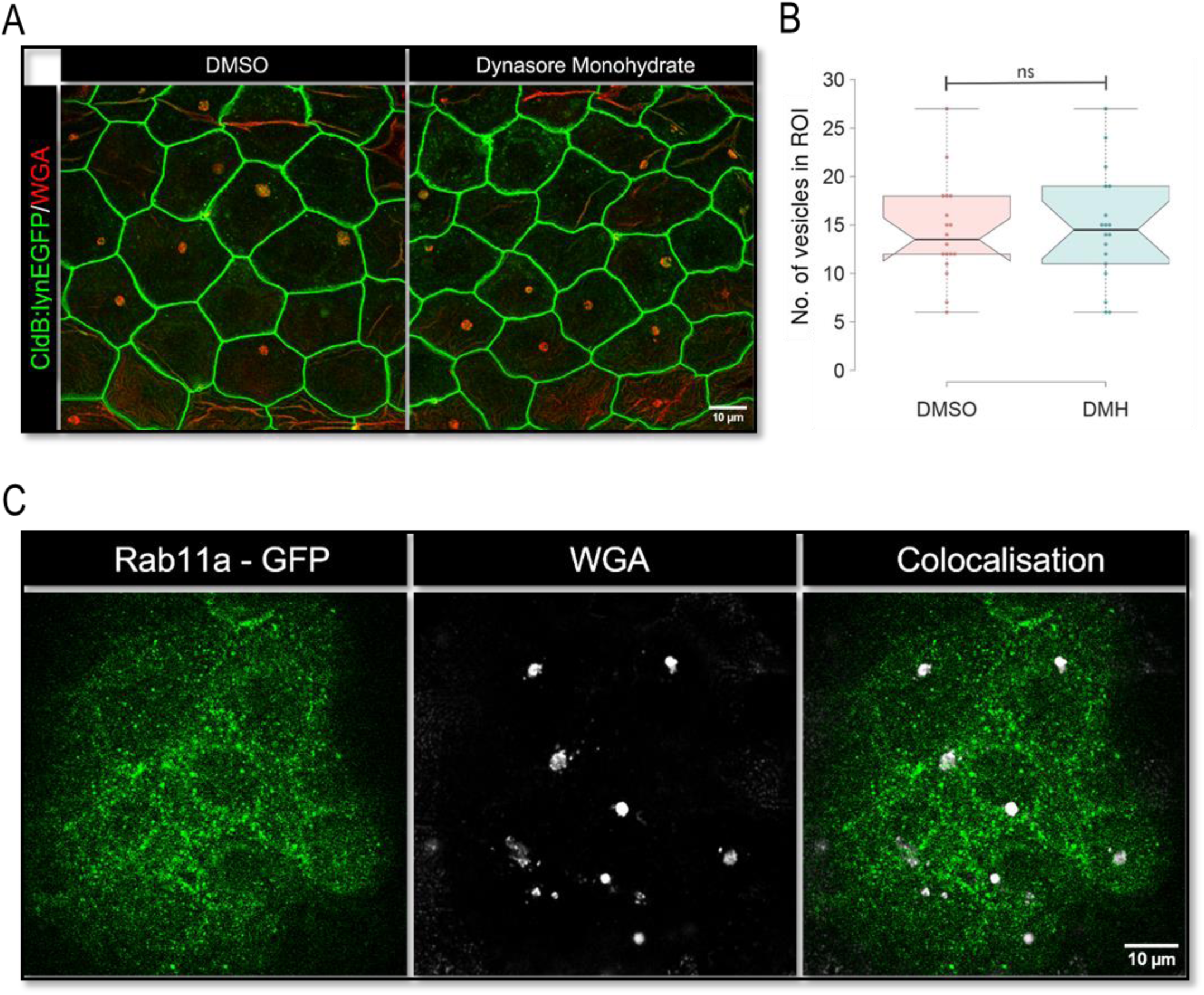
Formation of peridermal vesicles does not require Dynamin function and the vesicles are non-recycling endosomes. (A) Confocal scans of embryos treated with 30 μM Dynasore Monohydrate (DMH) from 48 to 72 hpf, followed by a 4-hour WGA pulse, illustrating the impact of dynamin inhibition on vesicle formation. (B) Notched box-plot of vesicle numbers per region of interest (ROI) from the embryos treated with DMH and vehicle control (DMSO); n = 18 embryos from three experimental sets; ns = not significant by a two-tailed t-test. (C) Live confocal images of peridermal cells from Tg(Rab11:EGFP) embryos pulsed with WGA marking macropinocytosis derived peridermal vesicles and Rab11a-GFP marking recycling endosomes; Scale bar: 10 μm.

**Supplementary Figure 3:**
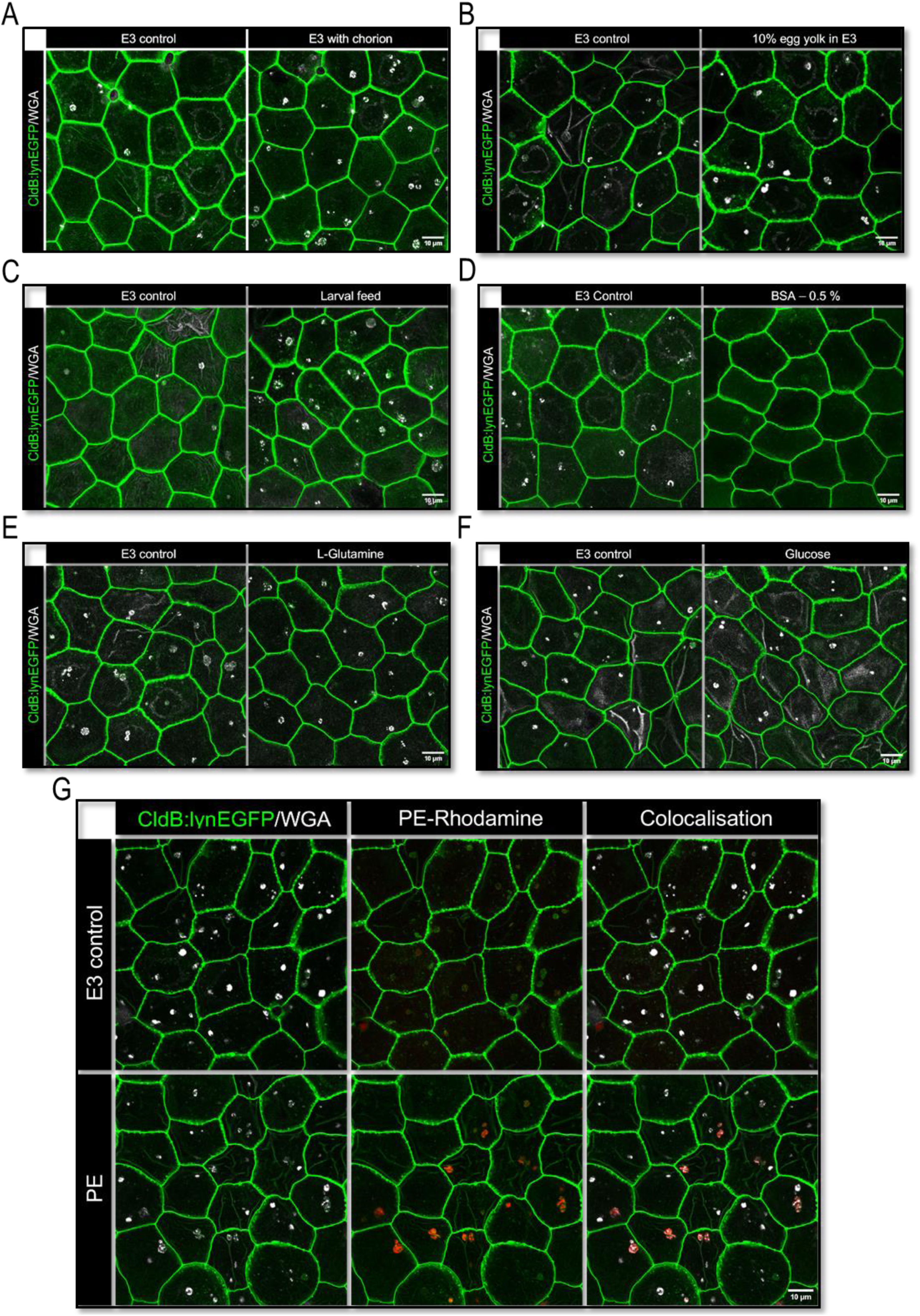
Effects of different supplementations on SNAC formation in zebrafish periderm. Confocal images of peridermal cells marked with lyn-EGFP at 72 hpf, following WGA pulse labelling (A-G) after 24h supplementation of E3 medium with (A) chorion, (B) 10% chicken egg yolk, (C) 4 mg/ml zebrafish larval feed, (D) 0.5% BSA, (E) 75 mM L-Glutamine, (F) 30 mM glucose and (G) 10 μg/ml Rhodamine-labelled phosphoethanolamine lipid.

**Supplementary Figure 4:**
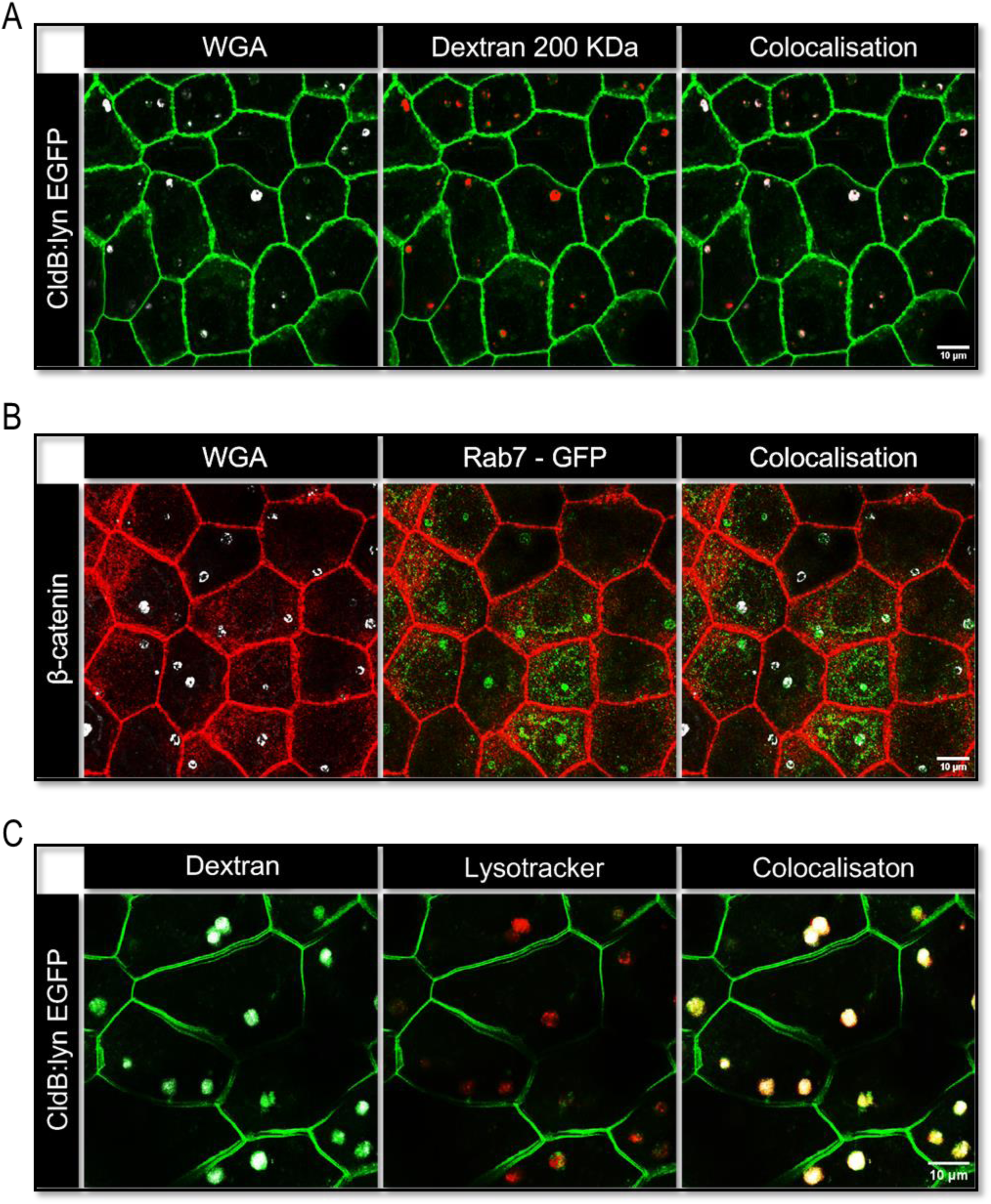
Characterization of SNACs in the Zebrafish Periderm. (A) Confocal micrographs of the periderm labelled with lyn-EGFP and pulsed with 200 kDa dextran and WGA at 72 hpf, following zebrafish yolk-stimulation at 48 hpf, showing macropinocytic uptake of dextran leads to the formation of SNACs. (B) Confocal imaging of 24h yolk stimulated peridermal cells in *Tg(Rab7:EGFP)* embryos immunostained for β-catenin showing late endosomes labelled with Rab7-EGFP and SNACs identified by a 4-hour WGA pulse at 72 hpf. (C) Confocal live images of Lysotracker and Dextran labelled peridermal cells following a 24h zebrafish embryo yolk stimulation at 48 hpf, revealing acidification of SNACs through co-labeling with Lysotracker and dextran during a 4-hour pulse at 72 hpf.

